# Evidence Graphs: Supporting Transparent and FAIR Computation, with Defeasible Reasoning on Data, Methods, and Results

**DOI:** 10.1101/2021.03.29.437561

**Authors:** Sadnan Al Manir, Justin Niestroy, Maxwell Adam Levinson, Timothy Clark

## Abstract

**Introduction:** Transparency of computation is a requirement for assessing the validity of computed results and research claims based upon them; and it is essential for access to, assessment, and reuse of computational components. These components may be subject to methodological or other challenges over time. While reference to archived software and/or data is increasingly common in publications, a single machine-interpretable, integrative representation of how results were derived, that supports defeasible reasoning, has been absent.

**Methods:** We developed the Evidence Graph Ontology, EVI, in OWL 2, with a set of inference rules, to provide deep representations of supporting and challenging evidence for computations, services, software, data, and results, across arbitrarily deep networks of computations, in connected or fully distinct processes. EVI integrates FAIR practices on data and software, with important concepts from provenance models, and argumentation theory. It extends PROV for additional expressiveness, with support for defeasible reasoning. EVI treats any computational result or component of evidence as a defeasible assertion, supported by a DAG of the computations, software, data, and agents that produced it.

**Results:** We have successfully deployed EVI for large-scale predictive analytics on clinical time-series data. Every result may reference its evidence graph as metadata, which can be extended when subsequent computations are executed.

**Discussion:** Evidence graphs support transparency and defeasible reasoning on results. They are first-class computational objects and reference the datasets and software from which they are derived. They support fully transparent computation, with challenge and support propagation. The EVI approach may be extended to include instruments, animal models, and critical experimental reagents.

## 1 Introduction

### 1.1 Motivation

It is now increasingly understood that dramatically enhanced capabilities for generating and analyzing very large datasets, with increasingly sophisticated methods, require systematic referencing of archived datasets and software as persistent first-class objects, with some machine-readable record of their provenance. There is now beginning to be an additional, and necessary, focus on software citation, and provenance. Principles and methods for achieving these goals have been defined and are in various stages of transition to practice in the scientific communications ecosystem [1–11].

Incorporating these practices has been advocated to improve verifiability, replicability, reproducibility, and reusability of computational results. The goal, which was established as a requirement at the dawn of modern science [12–15], is to make the process by which results and the claims they support are arrived at, transparent, and to allow the methods involved to be inspected – at least virtually – for adequacy, and if possible, reused, and improved upon in various applications.

We would like to do this for computations, using a clean, formal, and integrated approach, that does not require “boiling the ocean”; and in which artifacts such as provenance records, which may benefit a broad community of research, are generated principally as side-effects of normal computational work. We do not want to place unwanted burdens or requirements on researchers, with which they cannot realistically be expected to comply. Any system or method that generates such artifacts ought to have other attributes of significant value to researchers. In particular, we would like an ontology providing these features to provide useful functionality in a *digital commons* environment, where asynchronous reuse of results by various researchers occurs, and not necessarily as coherent workflows.

Of course, all published scientific results (including mathematical proofs) [16–19], are provisional. Results, methods, reasoning, computations, and interpretations may be challenged by others in the community, and frequently are [20, 21]. A major reason for methodological transparency is to support such reviews and challenges. Results in science are reviewed “by a jury of peers”, similarly to adversarial proceedings in law: a case is made, which may be argued for and against, based on evidence. Published “findings” or claims, and indeed computational results, are not facts. They are *defeasible assertions* [22], which rely upon a chain of evidence as warrants for belief. That makes them part of a chain of argumentation. We, therefore, treat computations and their provenance as defeasible arguments for provisional results.

### 1.2 Related Work

We previously undertook an analysis of the scientific communications life cycle to develop the Micropublications Ontology (MP) [23], which introduced a focus on chains of evidence and defeasible reasoning, and an emphasis on the nature of scientific claims as embedded in arguments. This approach was inspired by argumentation theory and by Greenberg’s detailed model of citation distortion [24, 25], which highlights empirical issues with citation chains in the scientific literature. It was also motivated by a perceived tendency amongst some computer scientists to take claims in the biomedical literature as “facts”, without subjecting them to further scrutiny.

Argumentation frameworks [26–34] and abstract dialectical frameworks [35, 36] are important sets of tools and concepts developed in the broader AI community, with an extensive literature. Bipolar Argumentation Frameworks (BAFs) as developed by Cayrol and others [31, 37, 38], allow both supporting and challenging arguments to be asserted and reasoned over in formal models. We discuss the relationship of our work to formal argumentation frameworks further in the Methods section.

The W3C PROV model [39–41] provides a well-thought-out and extensively tested set of core classes and properties, which may be used to document almost any computational provenance, in arbitrarily fine detail, and can serve as the basis for useful extensions. While PROV deals comprehensively and rigorously with core internal aspects of provenance, it does not engage explicitly with the role that provenance may play as evidence for computational results and interpretive claims. Nor does it conceptualize provenance as part of an argument for the validity of results, which may be countered, for example by later researchers finding bugs in code [42–45], flaws in datasets [46, 47], statistical errors [20], or fallacies in mathematical arguments [18].

Ontologies or schemas directly engaging the topic of experiments and experimental results include the Evidence and Claims Ontology (ECO) [48]; the Investigation, Study, Assay (ISA) model [49]; and the Ontology of Biomedical Investigations (OBI) [50]. All of these works are capable of representing machine-interpretable instances of scientific experiments within their proposed models and of offering limited provenance information. However, their focus is on characterizing individual experiments – principally in the wet lab – sometimes in exquisite detail, with over 4,000 classes in OBI, which make it best suited to use by highly trained specialist annotators. None of these models directly treat computational results, and none treat results as components of argumentation.

The ambitious Nanopublications model [51–54], was developed to standardize the form of research claims by recasting them as RDF triples, and aggregating holotypic claims with their paratypes, thus making the claims computable in some sense. What the nanopublications model currently lacks, is the ability to show evidential support in argumentation for the results it models.

Finally, there are related works that deal with computational results by packaging them up with their provenance records and other materials for citation or reference in metadata. These would include the Research Objects (RO) model [55–57] and its companion, RO-Crate [58]. The initial RO publication states as an explicit goal, replacing the “static PDF” form of scientific publication, and as such RO provides an integration framework across many related models involved in scientific specification, description, and communication. RO-Crate is a lightweight packaging initiative, or implementation realization, for RO and related material.

## 2 Methods

Cayrol and Lagasquie-Schiex’s work on BAFs supplied inspiration for our approach; which has, however, somewhat different semantics from their model, regarding the support relation, and the meaning of an argument.

In BAFs, arguments are opaque, and without internal structure. This extremely abstract treatment derives from Dung’s original presentation [33], enhanced to provide explicit bipolarity. In BAFs, if an argument A supports B, it agrees with B, similar to having an ally in a dispute. Therefore, an attack of C upon A also attacks B.

Our model, in contrast, treats support as it occurs in the scientific literature, as supporting evidence cited by the author of an argument. If C challenges A, a challenge by C on the supporting evidence for A cannot be inferred. However, as in Toulmin argumentation, and the work of Verheij [59–61], a challenge to A’s supporting evidence undercuts A – it reduces the *warrant for belief* in A.

Evidence graphs in our approach may be represented abstractly as:

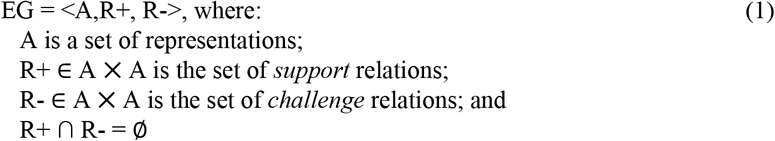

In our treatment, Representations are generalized forms of the various types of assertions and evidence found in the scientific literature. All Representations have provenance, if only to the extent that they have authorship. A Representation may be a statement, or a set of assertions, claims, or declarations, with provenance as its support.

The simplest form of provenance is the attribution of an assertion to an *Agent*. A declarative sentence is an argument of this simple form, where a statement’s attribution is its simplest supporting evidence. This corresponds to Aristotle’s view of argument as having a significant measure of support from ήθoς (ethos), the character of the speaker [62]; and relates to our common experience of tending to give more credence to statements from trusted, highly reputed sources. This notion can be extended to results produced by computational agents whose operations are well-validated and transparent.

A *Method* or *Material* representation in a publication, provided as provenance, constitutes a set of assertions about what was done to get a result. Software source code is a set of assertions of this form when used in a provenance description for a computation.

Source code has a dual nature, in that as a set of instructions, it is a collection of performatives [63], not evaluable as strictly true or false. We distinguish here between software as a set of instructions to a computer, which does something; and software provided as a description of what was done by the computer, i.e. “it ran this code”.

We have adapted these notions to computational provenance representation. We base our model on the following key ideas:

- All data, methods descriptions, and results are sets of defeasible assertions.
- The evidence for the correctness of any result is the record of its provenance.
- Subsequent research or discussion may challenge results, datasets, or methods.

The EVI ontology is a formal representation of the evidence for any result or claim as an EvidenceGraph, which unifies existing models for both software and data citation and supports machine-based defeasible reasoning.

We first introduced several of the concepts used in EVI in our previous work on Micropublications. EVI simplifies, revises, and adapts many features of the micropublications model to a purely computational digital commons environment, where evidence graphs may be generated by a computation service. The computation service we developed, described elsewhere [64], provides affordances to the user by greatly simplifying access to very large-scale data and underlying parallel computation and workflow services. At the same time, it produces and extends evidence graphs transparently, as a side effect.

## 3 Results

EVI is an extension of W3C PROV, based on argumentation theory, which enables defeasible reasoning about computations, their results, data, and software. It can be extended to incorporate non-computational evidence important to the results of a computation, for example, the specific instrument (manufacturer and catalog number) used to obtain a dataset or the reagent used in a binding assay.

Evidence Graphs are directed acyclic graphs, DAGs, produced by a service when a computation is run. They are first-class digital objects and may have their own persistent identifiers and be referenced as part of the metadata of any result. They may be arbitrarily deep. We model these using an OWL 2 vocabulary and set of rules [65, 66] which provide for propagation of support and challenge relations, with direct supports/challenges distinguished from indirect. A diagram of the classes and relations is provided in **Fig. 1**.

**Fig. 1.**
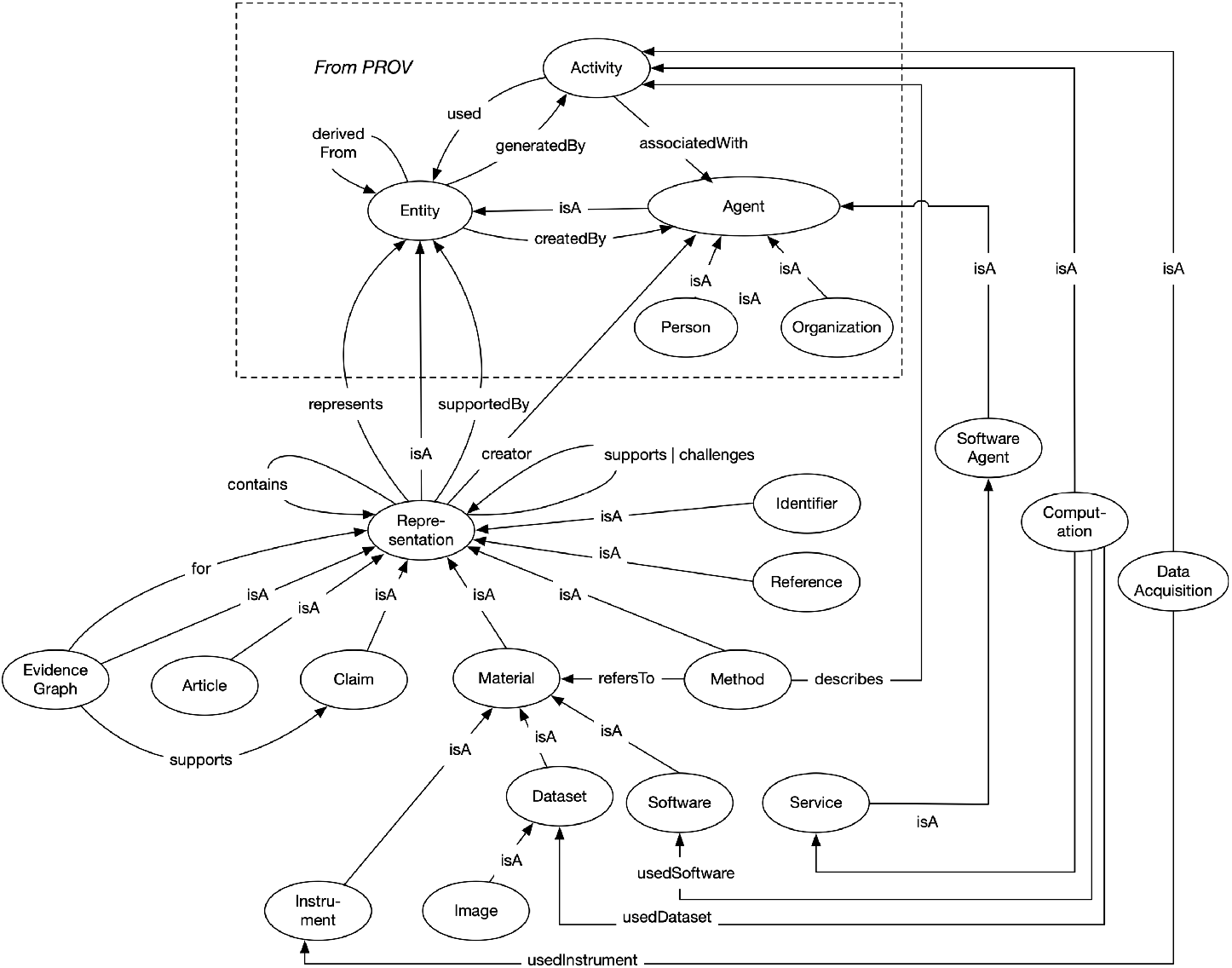
Classes and relations in the Evidence Graph Ontology, EVI. https://w3id.org/EVI

The classes within the dotted-line box in the figure are basic PROV classes: *Entity*, *Agent*, and *Activity*. All relations in the box are subproperties of PROV relations.

The argumentation relations *supports* and *challenges* are introduced on the new class, *Representation*, a subclass of *prov:Entity*. A *Representation* is a *prov:Entity* that *represents* another *Entity*. All digital objects in EVI are considered to be *Representations*; which may contain other *Representations*.

*Supports* and *challenges* are superproperties of *directlySupports* and *directlyChallenges* (not shown in the Figure). *Supports* is transitive, *directlySupports* is not. *DirectlySupports* is a superproperty of the relation *usedBy* (inverse of *prov:used*), and of the property *generates* (inverse of generatedBy) so that if a recorded *Activit*y D used a particular *Representation* C as input, and *generates* E as output, then C *directlySupports* D and D *directlySupports* E.

This is shown in Example 1 below, illustrating how the property chain rule evaluates distant support in the graph, where <s> stands for the *supports* property, and <dS> stands for *directlySupports*.

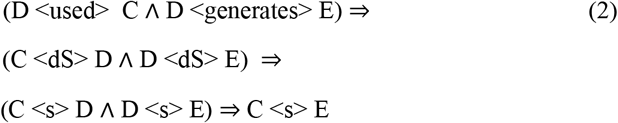

*Supports* and *directlySupports* properties are distinguished in this way because they serve different purposes in the model. The *directlySupports* property gives us our DAG, connecting the various Representations to form an evidence graph. It is simply a generalization over several PROV properties. The *supports* property is transitive and allows us to infer distant support relations. By analogy with genealogical trees, *directlySupports* equates roughly to *hasParent* relations; *supports* equates roughly to *hasAncestor* relations.

*Articles*, *Claims*, *Material* (as used in a typical “Materials and Methods” section), *Methods* (likewise), and *EvidenceGraphs* themselves, are *Representations*. A *Method* describes an *Activity* that may *referTo* a *Material*. The class *Material* is iteratively subclassed into *Instrument, Dataset, Image, Software*, and *Downloads*. *Downloads* are particular distributions of a *Dataset* or *Software*, representing e.g. different formats of a *Dataset*, in the same manner, this approach is used in the schema.org class *DataDownload*. An *Agent* in EVI may be a *SoftwareAgent*, *Person*, or *Organization*. A *Service usedBy* a *Computation* is a kind of *SoftwareAgent*. An *Activity* may be a *DataAcquisition* or a *Computation*.

Detailed formal definitions of all these terms, and equivalent schema.org [67] terms where they exist, are provided in the ontology description page at https://w3id.org, which has the latest updates. The OWL 2 vocabulary and its version history are on GitHub, here: *https://github.com/EvidenceGraph/EVI/blob/master/Ontology/versions/v0.2/evi.owl*, and the version we used for this article, is archived on Zenodo [68].

The diagram in **Fig. 1** omits detail relevant to rules and supports/challenges propagation, for pictorial clarity. The *supports* and *challenges* properties have subproperties *directlySupports*, *indirectlySupports*, *directlyChallenges*, and *indirectlyChallenges*, not shown in the figure. The *supports* property is propagated through the evidence graph via transitivity on *supports*.

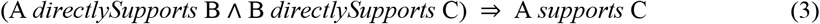

Challenges are propagated through a property chain on *challenges* and *supports*.

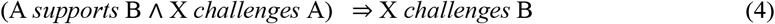

## 4 Discussion

In developing and validating our approach, we performed a very large-scale time series analysis of ten years of vital signs time series data from the Neonatal Intensive Care Unit (NICU) at the University of Virginia Hospital, collected on 5,957 infants. This extended proof-of-concept was a significant research study in its own right, in which NICU time series were analyzed using 80 different algorithms in 11 mathematical families from diverse domains [69]. Computations were performed over a period of 15 months, in various stages, using the FAIRSCAPE microservices framework [64], producing an evidence graph of 17,996 nodes [70]. This evidence graph is queryable from a Stardog™ quad store via one of the microservices and is also deposited in the University of Virginia’s Dataverse with a *prov:wasDerivedFrom* property associating it with the underlying dataset, which is also in Dataverse [71].

Whether using our FAIRSCAPE framework or some other approach, EVI statements are intended to be transparently generated by a *computation service*, similar to how ordinary PROV statements are generated by workflow engines. Thus, evidence graphs become side effects of doing useful computations. In our case, these computations are run within a FAIR digital commons environment.

**Fig. 2** illustrates a section of this graph for one of the 5,957 subjects in the NICU analysis. In our microservices framework the EvidenceGraph service stores and retrieves the graphs in a StardogTM RDF quad store, which also performs the inferencing. The PATH query, an extension to SPARQL, generates an *EvidenceGraph* DAG from the root object, using the *directlySupportedBy* abstraction (not shown here) to structure the graph and the *supports* superproperty of its transitive inverse, *supportedBy*, to infer distant evidential support.

**Fig. 2.**
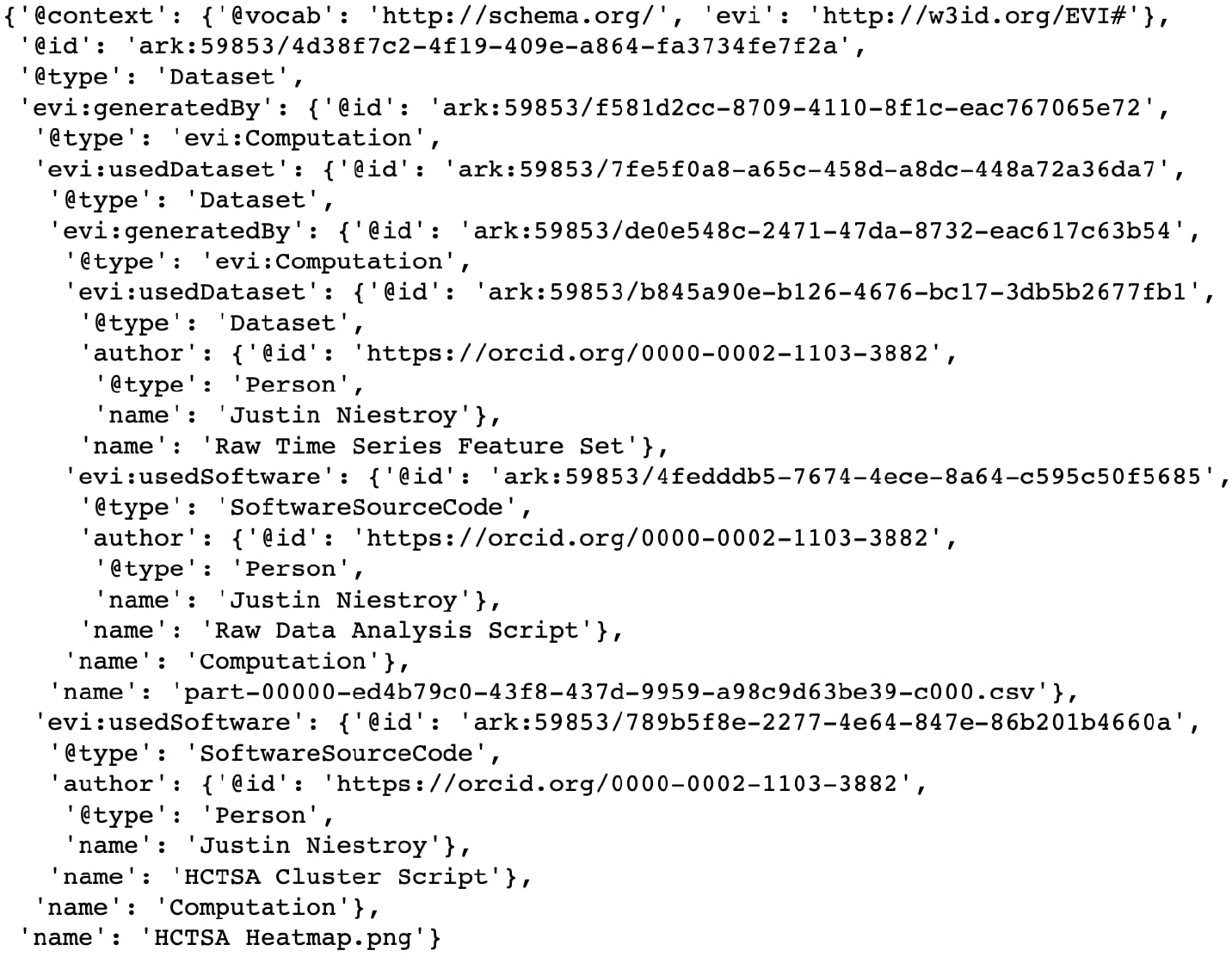
JSON-LD of a portion of the evidence graph for one of the 5,997 infant subjects from clustering step in comparative time-series analysis, adapted from [64].

As can be seen in the example, every node in the graph has a persistent identifier, based on the ARK system [72] in our implementation. This approach provides the important unification of all available digital evidence for (and potentially, against) a result, by supporting persistent resolution to the cited objects.

One could ask, in a practical sense, if a challenge is made to some leaf node deep in the graph, should it always invalidate the root assertion? The answer to this is that challenges do not invalidate, they present an opposing view. They ultimately require human judgment as to their validity and strength. We do, however, wish to know about them. This becomes important in digital commons environments, and in, for example, metaanalyses reusing prior results. It can have a further impact if computational results are properly referenced and tied to textual claims in citation networks, such as those explored by Greenberg [24] and others.

We believe that EVI provides an important generalization of provenance as evidence for correctness, which can be further extended beyond computation to include the other types of evidence presented in scientific publications, for example by including identifiers such as RRIDs [73] of important experimental reagents and animal models in the protocols from which the data was derived. Our end goal is to be able to provide, with the metadata of any datasets we store in an archive, a link to the corresponding evidence graph. With this practice, any result presented in a publication, with appropriate data citation, ought to be resolvable to its entire evidence graph, and transitively closed to its components. Future research directions include extended support for packaging evidence graphs; support for extended descriptive metadata; service integration with data and software archives; and continued alignment with other initiatives in this space. As a component of the FAIRSCAPE digital commons framework, we plan for EVI to continue its evolution with direct input from computational users.

## Information sharing statement

- The EVI ontology OWL2 vocabulary is available at https://w3id.org/EVI# under MIT license.

## Acknowledgements

We thank Chris Baker (University of New Brunswick), Carole Goble (University of Manchester), and John Kunze (California Digital Library) for helpful discussions. This work was supported in part by the U.S. National Institutes of Health, grant NIH 1U01HG009452; and by a grant from the Coulter Foundation.

